# 100+ years of bird survey data reveal changes in functional fingerprints indexing ecosystem health of a tropical montane forest through time

**DOI:** 10.1101/2020.06.30.180950

**Authors:** Camila Gómez, Elkin A. Tenorio, Carlos Daniel Cadena

## Abstract

Ecologically relevant traits of organisms inhabiting an ecosystem determine its functional fingerprint. Quantifying changes in the shape, volume and shifts in the position of functional fingerprints can provide information about the effects of diversity loss or gain through time, and is a promising means to monitor ecological integrity. This, however, is seldom possible owing to limitations in historical surveys and lack of data on organismal traits, particularly in diverse tropical regions. Using detailed bird surveys from four time periods across more than one century and morphological traits of 233 species, we quantified changes in the avian functional fingerprint of a tropical montane forest site in the Andes of Colombia. We found that 79% of the variation in functional space, regardless of time period, was described by three major axes summarizing body size, dispersal ability, and habitat breadth. Changes in species composition caused significant alterations of the functional fingerprint of the assemblage, with 35 – 60% reductions in functional richness and dispersion. Owing to species extirpations and to novel additions to the assemblage, functional space is currently smaller and at least 11% different to what it was a century ago, with fewer large-sized species, more good dispersers, and fewer habitat specialists. Extirpated species had high values of functional uniqueness and distinctiveness, resulting in large reductions of functional richness and dispersion after their loss, implying potentially important consequences for ecosystem functioning. Conservation efforts aimed at maintaining ecosystem function must move beyond maintaining species numbers to designing strategies for the maintenance of ecological function by identifying and conserving species with traits conferring high vulnerability.

## Introduction

Determining how changes in the natural world affect ecosystems and the biodiversity within them is essential, especially if we are to maintain ecosystem function and restore degraded habitats to regain services they once provided (Cadotte et al. 2011; Dirzo et al. 2014). First, however, we must understand what defines healthy ecosystems in terms of function, and how ecosystems change naturally or in response to human intervention. Functional traits, defined as ecological, morphological or behavioral traits influencing fitness and survival of organisms (Violle et al. 2007), mediate ecosystem processes, and determine the responses of populations to environmental conditions, biotic interactions and nutrient cycling in ecological assemblages (Bregman et al. 2016; Funk et al. 2017). Therefore, measures of diversity in functions provide information about ecosystem health because higher functional diversity is associated with greater plasticity, productivity, and resilience when facing disturbances or climatic fluctuations (Mason et al. 2005; Cadotte et al. 2011). Examining changes in functional diversity within assemblages known to have changed in species composition through time offers an opportunity to understand the consequences that impacts on biodiversity have on the functions and services it provides.

Measures of functional diversity are often quantified as a multidimensional volume, where each species occupies a position depending on its similarity to others. In this volume, species with similar trait values (i.e. functionally redundant species) are located in the center close to other species, whereas functionally distinctive species are far from the rest in the periphery (Mason et al. 2005; Ricklefs 2012; Kuebbing et al. 2018). The shape of the multi-dimensional trait space formed by organisms inhabiting an ecosystem hence determines a unique “functional fingerprint”. Quantifying changes in the shape, volume and shifts in the position of functional fingerprints can therefore provide information about the effects of diversity loss or gain, and is a promising means to monitor ecological integrity (Carmona et al. 2016; Pigot et al. 2016; Cooke et al. 2019b).

In response to environmental changes, the functional fingerprint of an assemblage may remain unchanged, shift in different directions, or change in shape and volume. For instance, if an assemblage loses or gains species far from others in functional space, then its functional richness (i.e. the total functional space it occupies) would change more drastically than if it lost or gained species occupying similar areas of trait space relative to other members of the assemblage (Carmona et al. 2016; Grenié et al. 2017; Kuebbing et al. 2018). Similarly, whether lost or gained species are located close or far from the centroid of trait space will determine the effect of changes on functional dispersion (i.e. the mean distance of each species to the centroid of trait space; Laliberte & Legendre 2010), with extirpations of species in the periphery of functional volume causing greater decreases in functional dispersion (Laliberte & Legendre 2010). Finally, changes in abundance of species in an assemblage can also impact its functional dispersion by shifting the center of gravity towards the most abundant traits (Laliberte & Legendre 2010). Changes in functional shape or volume would therefore depend on characteristics of the species changing in abundance relative to the rest of the assemblage (Carmona et al. 2016).

Birds are good indicators of ecosystem health because they rapidly respond to environmental changes, and because variations in bird assemblages can be readily assessed by human observers (e.g. Gregory & van Strien 2010; Bregman et al. 2016). Furthermore, the link between avian functional traits and ecosystem processes is well documented (Maglianesi et al. 2015; Ikin et al. 2019; Pigot et al. 2020). Although many studies have documented changes in bird assemblages through time as a response to habitat or climate change (e.g. Renjifo 1999; Robinson 1999; Freeman et al. 2018; Rosenberg et al. 2019), determining whether ecosystem health is affected by changes in species diversity and abundance requires assessing whether changes extend to functional spaces.

Studies of the temporal dynamics of functional diversity within ecological assemblages indicate that extinctions and colonizations of species are not random in terms of their functional traits (Petchey et al. 2007), and changes in functional space characterize succession in ecological assemblages (Purschke et al. 2013). Responses to habitat and landscape changes also affect functional groups differentially and produce changes in interspecific interactions which affect ecosystem function at different scales (Chalmandrier et al. 2015; Jarzyna & Jetz 2018; Stouffer 2020). Furthermore, changes in functional diversity after habitat fragmentation may homogenize traits within assemblages, implying loss of unique functions (Clavel et al. 2011; Jarzyna & Jetz 2017). However, studies quantifying long-term changes in the multidimensional functional fingerprint of tropical assemblages are lacking.

The avifauna of San Antonio, a montane forest locality in the Andes of Colombia, poses an unprecedented opportunity to examine shifts in functional fingerprints through time in a highly diverse tropical ecosystem, and thereby to infer how species extirpations and recolonizations may have influenced ecosystem health. San Antonio was first surveyed by naturalists in the 1910’s (Chapman 1917). Since then, exhaustive resurveys conducted in the 1950’s, 1990’s and 2000’s have allowed researchers to document substantial changes in composition of the bird assemblage over more than a century of landscape change (Kattan et al. 1994; Palacio et al. 2019). We combined morphological and ecological data for the complete bird assemblage of San Antonio, comprising 233 species, to test the hypothesis that functional fingerprints changed as a result of gains, losses and changes in abundance of functionally distinctive species. We expected functional richness and dispersion to have decreased between the early 1910’s and the 1990’s when several species extirpations occurred (Kattan et al. 1994), and to have increased following recovery of some species and colonization by formerly absent species in the 2000’s (Palacio et al. 2019). We also evaluated whether colonization by novel species caused changes in the functional fingerprint of the San Antonio assemblage, filling areas of functional space not occupied by the set of species coexisting in the area in the 1910’s. If extirpated species were functionally unique, then this system may have lost functions provided by those species and potentially gained others from the novel colonizers. Alternatively, if extirpated species or novel colonizers were functionally redundant, then the overall functionality of the system may not have changed significantly.

## Methods

### Study site

The *Cerro de San Antonio* is a mid-elevation mountain ridge (1700 - 2200 m), located in Colombia’s Western Andes (3.4960 N, −76.6305 W) approximately 15 km west of Cali, Valle del Cauca (Kattan et al. 1994; Palacio et al. 2019). This region (covering ~7000 ha) originally harbored extensive tropical montane and cloud forests, which suffered widespread fragmentation from the 1930’s to the 1960’s resulting in a ~46% reduction in forest cover (Kattan et al. 1994; Palacio et al. 2019). Since then, the remaining matrix of forest fragments, small farms and country houses has remained relatively stable, with an estimated ~10% increase in forest cover between 1995 and 2016 (Palacio et al. 2019).

### Historical and contemporary bird survey data

We analyzed data based on published bird lists from San Antonio compiled by Palacio et al. (2019), which comprise a total of 233 species detected across surveys conducted between 1907 and 2016 (Appendix S1). We divided data into 4 periods: (1) the 1910’s, corresponding to surveys led by Frank M. Chapman and Mervin G. Palmer; (2) the 1950’s, by Alden H. Miller; (3) the 1990’s, by Gustavo Kattan et al.; and (4) the 2000’s, by Ruben Palacio et al. Detailed methodologies for these surveys can be found in the original publications (Chapman 1917; Miller 1963; Kattan et al. 1994; Palacio et al. 2019) and together they comprise an accurate representation of the avifauna of San Antonio and its compositional changes over 100+ years. For details on changes in the avifauna during these time periods see Appendix S3.

Surveys in the 1910’s consisted exclusively of collecting expeditions (Chapman 1917) but from the 1950’s onwards, surveys combined standardized observations with collecting, and the most recent surveys in the 2000’s also integrated citizen-science data (Kattan et al. 1994; Palacio et al. 2019). Methodological differences between surveys imply that we have strong certainty about species extirpations occurring between time periods but are less certain about novel species colonizations because the older surveys were more likely to have missed species which were actually present but not identified or collected due to limited knowledge of bird songs, lack of good binoculars, etc.. To address this issue, we analyzed our data considering two scenarios. The first assumes that all surveys accurately describe the avifauna present and that any records of species not detected in prior surveys of San Antonio represent real colonizations. The second, more conservative scenario, assumes that all the ‘novel species’, i.e. those detected for the first time after the 1910’s, were false absences and so we added them to the 1910’s list. Therefore, under Scenario 2 any changes in the avifauna consisted exclusively of extirpated species or re-colonization of previously extirpated species. What actually occurred likely lies somewhere in the middle of our two scenarios.

### Functional traits

For all 233 species, we compiled information on 9 traits describing functional morphospace and ecological strategies (Cooke et al. 2019b; Habel et al. 2019; Pigot et al. 2020; Sheard et al. 2020): body mass, bill length, bill width, wing chord, tail length, tarsus length, hand-wing index (i.e. a measure of wing shape), habitat breadth and generation time (BirdLife International 2018; Appendix S1). We also made sure most of the traits considered had low values of correlation within our assemblage (72% of traits have correlation coefficients < 0.6, supplementary Fig. S3.1 in Appendix S3; Cadotte et al., 2011). For additional details on functional trait compilation see Appendix S3.

### Estimation of functional diversity metrics and temporal comparisons

Body mass was log10 transformed, and all traits were scaled and centered to have zero mean and unit variance (Cadotte et al. 2011; Carmona et al. 2016; Cooke et al. 2019a). To obtain a reduced set of uncorrelated variables explaining variation in functional traits, we ran a principal components analysis using data from all species (R Development Core Team 2019). Out of nine principal components, the first three explained 78% of the variation in functional trait values. Loadings suggested that PC1 described mostly variation in body size (55%), PC2 was related to dispersal ability through the hand-wing index (13%) and PC3 described habitat breadth (11%; Appendix S3). The first 3 PC scores were then used as axes to compare functional spaces among time periods and groups of species (Carmona et al. 2016; Cooke et al. 2019b).

We adopted methods from Cooke et al. (2019) to construct functional spaces and evaluate temporal shifts in the San Antonio avifauna since the 1910’s. Two and three-dimensional trait spaces are constructed by comparing different combinations of trait-space values, in this case PC1, PC2 and PC3 (Cooke et al., 2019). Multivariate kernel density estimates (Duong 2019) were used to calculate the 0.5, 0.95 and 0.99 probability contours for trait spaces (Cooke et al. 2019b). Scores from PC’s comprising ~95% of variance were used to construct a trait hypervolume to visualize temporal shifts in functional space and to calculate changes in volume and overlap between time periods. The hypervolume was constructed following Cooke et al. (2019) using a ‘one-class support vector machine (SVM)’ method (Blonder & Harris 2019).

Historical and contemporary relative abundances of species from San Antonio were extracted from Palacio et al. (2019). We assigned a value between 0 and 1 to each abundance category where 0 = absent or extirpated, 0.2 = rare, 0.4 = uncommon, 0.6 = fairly common, 0.8 = common, and 1 = abundant. We constructed a period-by-species matrix with these relative abundance values, and a species-by-trait matrix with the standardized values of our 9 functional traits. We then estimated functional richness and dispersion using function ‘*dbFD*’ in R package ‘*FD*’ (Villeger et al. 2008; Laliberte & Legendre 2010; Laliberté et al. 2014). Functional richness (*FRic*) was estimated as the volume of the minimum convex hull (Villeger et al. 2008). For ease of interpretation, the functional richness of each time period was standardized by the global *FRic*, thus constraining values to range between 0 and 1 (Laliberté et al. 2014). Functional dispersion (*FDis*) is the mean distance of each species to the centroid of trait space, weighted by its relative abundance (Laliberte & Legendre 2010). *FDis* increases as the most abundant trait values are further from the center of gravity of trait space (Laliberté et al. 2014). The relative abundance categories for the historical datasets may be inaccurate because they do not account for collecting bias, i.e. an uncommon species in the collection might have been common in the assemblage. We therefore also estimated *FDis* without considering abundances.

To evaluate whether functional metrics for each time period were different from what might be expected through random processes, we compared observed values to those generated by null models. We randomly reordered the species-identity column of the abundance matrix 999 times (thus breaking associations between traits and species’ identities), and then recalculated *FRic*, and *FDis* for each random assemblage (Van de Perre et al. 2020). To assess the magnitude of the difference between observed and null values of *FRic* and *FDis*, we estimated standard effect sizes as: 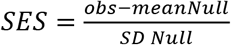 (Van de Perre et al. 2020) and assessed significance with *P* values as: 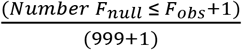 (Cooke et al. 2019a).

### Functional characteristics of extirpated species and novel colonizers

To describe functional differences among 1. extirpated species, 2. novel additions to the assemblage as of the 2000’s, and 3. shared species maintained through time, we constructed trait probability density curves (TPD), using the first three PC axes as traits, and the aforementioned species groups as ecological units for comparison in package ‘TPD’ (Carmona et al. 2016, 2019). We used a two-sample Kolmogorov-Smirnov test to assess whether there were differences between TPD’s of extirpated species and novel additions to the assemblage.

Additionally, to evaluate whether extirpated species and those establishing new populations in San Antonio were functionally redundant or unique, we calculated functional distinctiveness and uniqueness using package ‘funrar’ in R (Grenié et al. 2017; Pimiento et al. 2020). Functional distinctiveness (*D_i_*) measures how uncommon a species’ trait value is compared to other species in the assemblage, weighted by the species’ relative abundance, and has values between 0 (high functional redundancy), and 1 (low functional redundancy; Grenié et al. 2017). Functional uniqueness (*U_i_*) is the distance of each species to its nearest neighbor in the assemblage (Grenié et al. 2017). The closer *U_i_* is to 1, the further species *i* is to its closest neighbor (Grenié et al. 2017). We ran permutation tests to evaluate whether extirpated species and new additions to the assemblage were more functionally unique or distinctive than expected by chance (Pimiento et al. 2020), and carried out pairwise comparisons to assess differences between time periods (Cooke et al. 2019a).

## Results

Under both assumed scenarios (1 – species were both extirpated and colonized the assemblage; and 2 – there were no novel colonizers), the center of abundance of two-dimensional morpho-spaces has remained in the same position, but both the shape and extent of trait space have changed over 100+ years (Fig. 1 and supplementary Fig. S3.2). Extirpated species (both scenarios), as well as new colonizers (Scenario 1), are spread over functional space, but those located towards the periphery have been responsible for the most noticeable shape changes in functional space (Fig. 1 and supplementary Fig. S3.2).

**Figure 1.**
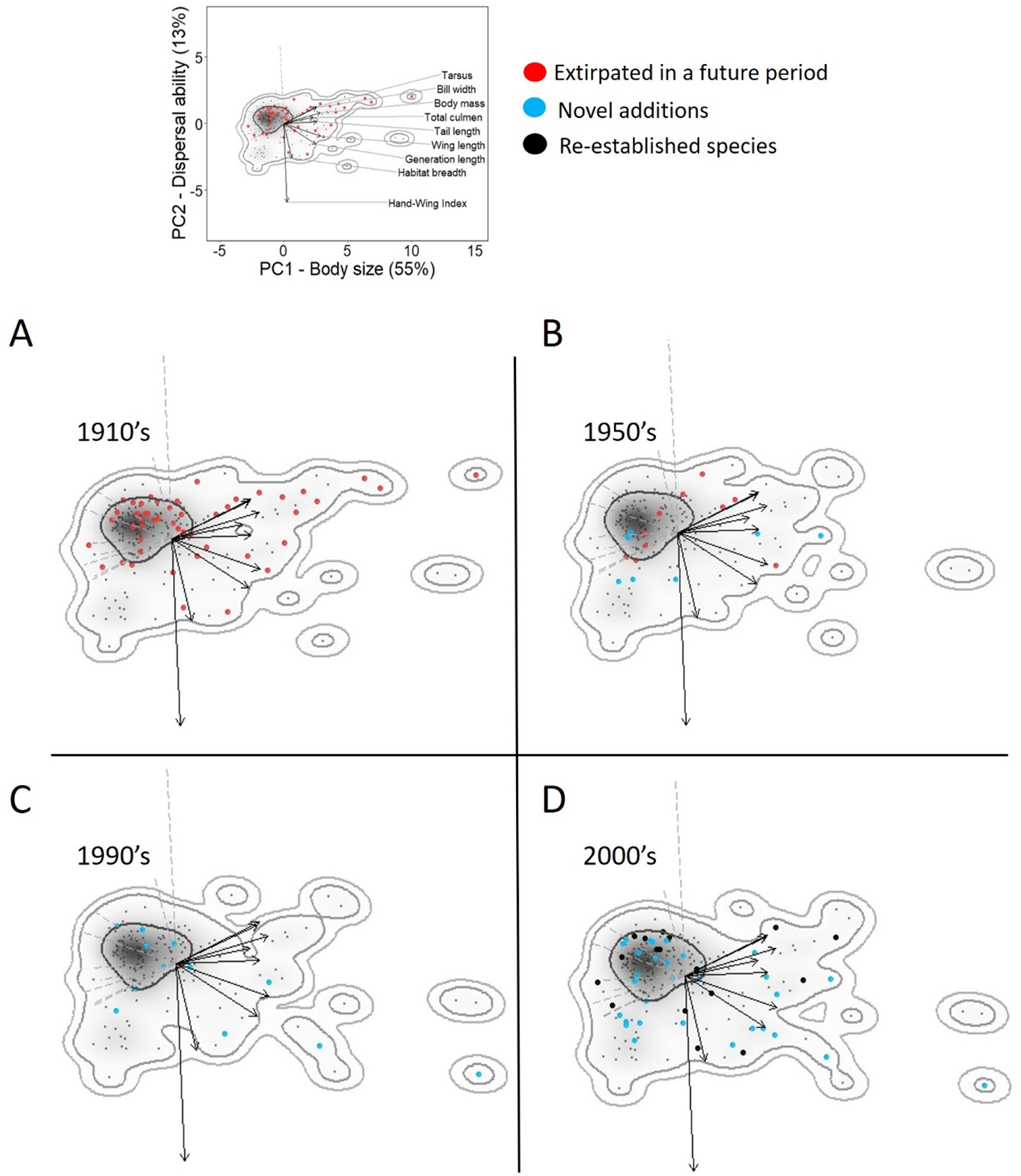
Species extirpations, novel additions and re-establishments have caused changes in the shape and extent of the functional space of the San Antonio avian assemblage in 100+ years, but the centroid of trait space (gray shading) has not shifted. Two-dimensional functional space is represented by PC scores of functional traits during 4 time periods (**A-D**). PC1 reflects largely variation in body size whereas PC2 correlates with dispersal ability of birds and habitat breadth. Small dots are species present in each time period, with larger red dots representing species which became extirpated in a future period and larger blue dots representing new additions which were absent in previous periods. In **D**, the larger black dots are species which re-established populations after being extirpated. Arrows are scaled to represent the loadings and direction of each trait in functional space and the insert above shows the scale and the traits represented by each arrow. Gray shading represents the kernel density estimates for each time period and curved lines show the 50, 95 and 99% probability contours.

Changes in functional space were also evident in trait density curves comparing extirpated species, novel additions, and shared species through time. Trait density curves for Scenario 1 showed that the San Antonio assemblage has seen a shift towards smaller-sized birds (k-s test D = 0.34, *P* = 0.03, Fig. 2A), and towards species with higher dispersal ability (k-s test D = 0.48, P = 0.0008, Fig. 2B) and wider habitat breadths (i.e. fewer habitat specialists; k-s test D = 0.44, *P* = 0.003, Fig. 2C). These shifts resulted in 34% of the functional hypervolume present in the 1910’s being absent from the 2000’s. In turn, 11% of the volume occupied by the 2000’s assemblage was not occupied in the 1910’s, and 55% has remained constant (Figure 2D). For Scenario 2, trait density curves for body size and dispersal ability showed similar shifts to those of Scenario 1 with the assemblage loosing larger-sized birds (k-s test D = 0.39, *P* = 0.0003) and some species with relatively low dispersal ability (k-s test D = 0.43, *P* = 0.000), but showing no significant shifts in habitat breadths of species (i.e. PC3; k-s test D = 0.24, *P* = 0.08) (supplementary Fig. S3.3). Under Scenario 2, 29% of the trait hypervolume from the 1910’s was lost, whereas only 1% of the hypervolume was unique to the 2000’s and 70% remained constant across periods.

**Figure 2.**
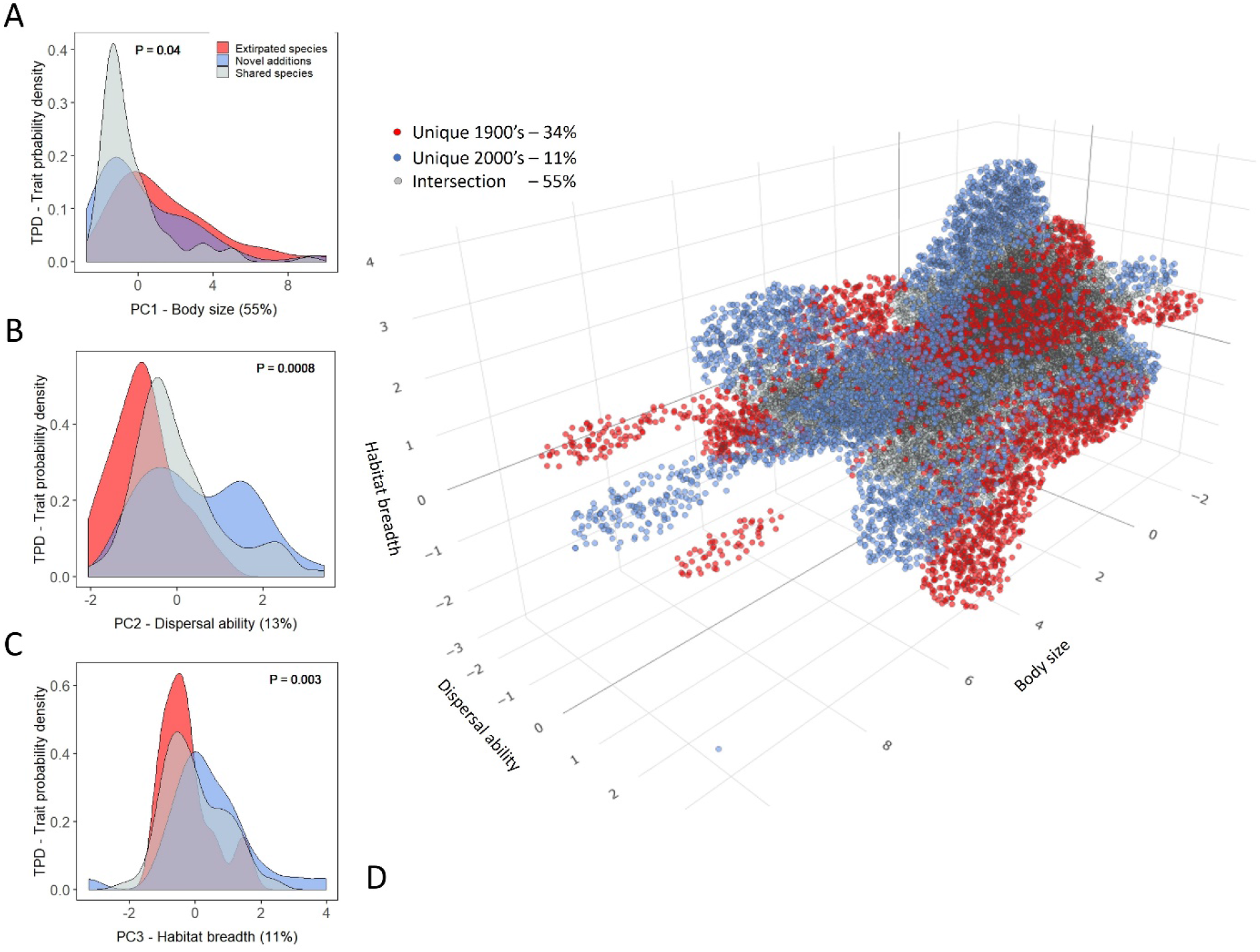
Probability density curves of extirpated species and novel additions to the avifauna between 1910’s and 2000’s (Scenario 1), show that San Antonio’s functional fingerprint has shifted towards **A.** Smaller birds, **B.** Species with higher dispersal ability and **C.** Species with wider habitat breadths. *P* values were obtained through Kolmogorov-Smirnov tests comparing the distributions of extirpated and novel species. **D.** An overlap of the 9-dimension hypervolumes of the 1900’s and 2000’s suggests that 35% of the volume occupied in 1910’s (in red) has been lost and replaced by 11% of new functional space (in blue) provided by the novel additions to the San Antonio assemblage during the 2000’s. 55% of the functional hypervolume has remained constant in time. Only the first 3 dimensions are used for visualization.

Changes in the hypervolume of all traits were comparable to measures of functional richness (*FRic*) using the minimum convex hull. Under Scenario 1, between the 1910’s and 1950’s, functional richness decreased by ~36%, and it recovered ~22% between the 1990’s and 2000’s (Table 1, Figure 3A). Under Scenario 2, there was a ~66 % decrease in functional richness between the 1910’s and 1990’s (Table 1) and then a recovery of 12% during the 2000’s (Table 1). Functional dispersion (*FDis*) followed a similar pattern in both scenarios (Table 1) with a decrease of ~8% between 1910’s and 1950’s and an increase of ~3% between the 1990’s and 2000’s (Table 1, Fig. 3B). The difference between values of *FDis* estimated with and withought weighting for abundance were negligible (mean difference = 0.02, range = 0.003 – 0.05). All observed values of *FRic* and *FDis* were lower than expected by chance, with negative standard effect sizes ranging from −0.87 to −2.81 (Fig. 3C).

**Figure 3.**
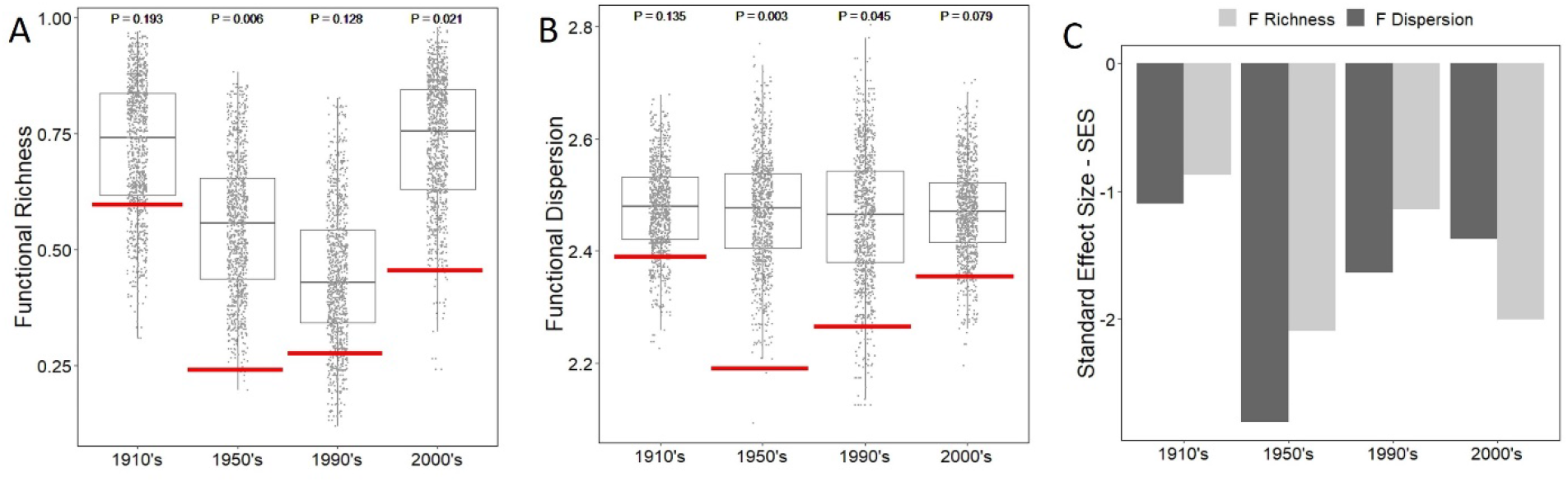
Functional richness (**A**), and dispersion (**B**) decreased significantly between the 1910’s and the 1950’s, appeared to increase slightly thereafter until the 1990’s and then increased significantly until the 2000’s. Red lines show the empirical values of functional richness and dispersion estimated for each time period, and boxplots show the distribution of these values generated by 999 iterations of a null model with randomized species identities. Deviance from random expectation, measured as standard effect sizes (**C**) were all negative suggesting that the assemblages at different time periods have lower than expected values of functional richness and dispersion, i.e. extirpated species were more functionally unique than expected by chance.

**Table 1.** Values of functional richness and functional dispersion estimated for two possible Scenarios of changes in species assemblages during 100 years in San Antonio – Colombia.

| Period | Functional Richness |  | Functional Dispersion |  |
| --- | --- | --- | --- | --- |
|  | Scenario 1<sup>a</sup> | Scenario 2<sup>b</sup> | Scenario 1 | Scenario 2 |
| 1910's | 0.60 | 1.00 | 2.39 | 2.45 |
| 1950's | 0.24 (- 36%)<sup>c</sup> | 0.41 (- 59%) | 2.19 (- 8%) | 2.28 (- 7%) |
| 1990's | 0.28 (+ 4%) | 0.34 (- 7%) | 2.27 (+ 3%) | 2.32 (+ 2%) |
| 2000's | 0.46 (+ 18%) | 0.46 (+ 12%) | 2.35 (+ 3%) | 2.35 (+ 2%) |

<sup>a</sup> Scenario 1 assumes there were novel species which joined the assemblage as well as extirpations and re-colonizations.
<sup>b</sup> Scenario 2 assumes there were no novel species, just extirpations and re-colonizations.
<sup>c</sup> Numbers are estimates for each time period and in parenthesis the equivalent change from the previous period as a percentage.

Finally, we found that the San Antonio assemblage has lost a higher number of functionally unique and distinctive species than would have been expected by chance (Figs. 4A-B), and pairwise comparisons between groups of species showed significant differences in functional distinctiveness and uniqueness between extirpated species and species remaining in the assemblage across time periods. For example, large frugivores, a group especially sensitive to extirpation, were among the highest ranked in functional distinctiveness and uniqueness (Fig. 4. and Appendix S3). Nine out of 32 novel additions to the assemblage (Scenario 1) also had relatively high values of functional distinctiveness and uniqueness, with the Ornate Hawk-Eagle (*Spizaetus ornatus*), which was first recorded during the 1990’s, having the highest value (Fig. 4).

**Figure 4.**
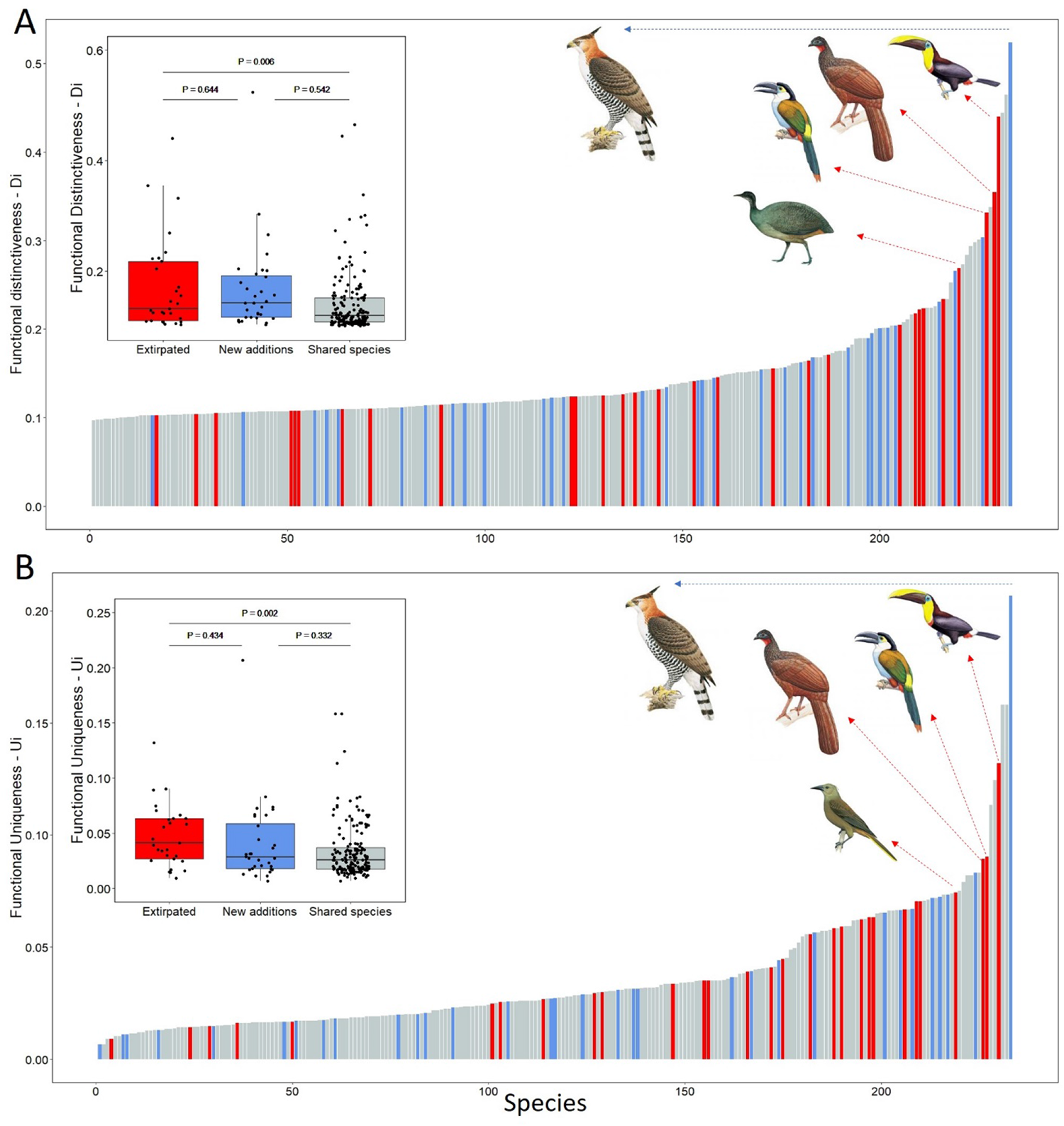
**Figure 4.** Both extirpated species and novel additions to the San Antonio assemblage had higher functional distinctiveness (**A**. *P_ext_* = 0.009, *P_new_* = 0.002) and uniqueness (**B**. *P_ext_* = 0.002, *P_new_* = 0.017), than expected by chance. Differences between pairs of groups were only significant between extirpated and shared species. The highest ranked extirpated species are all large frugivores, but the species with the highest rank is the Ornate Hawk-eagle, a novel addition which appeared in the 1990’s in San Antonio.

## Discussion

We showed that changes in species composition caused significant shifts in the functional fingerprint of a montane bird assemblage from the western Andes of Colombia. Due mostly to species extirpations, functional space in the 2000’s was considerably smaller and at least 11% different to what it was 100 years before, containing fewer large-sized species, more species with high dispersal ability, and fewer habitat specialists. However, the core of functional space, which holds most of the species (~65%), has remained in the same position, suggesting a portion of the system’s original functionality remains intact. On average, extirpated species had higher values of functional distinctiveness and uniqueness relative to the whole assemblage, and groups known to be globally prone to extirpation, such as large frugivores, were ranked high in uniqueness and distinctiveness. Therefore, despite the stability of the core functional space, important ecological functions performed by relatively few species may have been disproportionately affected by changes in assemblage composition.

We detected substantial (i.e. 30 – 60%) declines in functional richness in the San Antonio assemblage over 100 years. These values are high given that declines in functional richness in the 11-25% range are considered to cause substantial loss of functionality in highly diverse ocean and forest assemblages (Mcwilliam et al. 2018; Newbold et al. 2020; Pimiento et al. 2020), and that some systems with 40% difference in tree cover, such as agriculture vs old growth forest, differ by only ~5% in the functional richness of their bird assemblages (Ikin et al. 2019). Furthermore, we found that extirpated species had higher than expected values of functional uniqueness and distinctiveness, and therefore their loss was mostly responsible for the decreasing trend in functional diversity over 100 years. The joint substantial reduction in functional diversity and loss of unique species suggests that the San Antonio assemblage has likely lost important ecological functions.

Our approach can enable conservation managers to move beyond simple quantifications of losses and gains through time to identify areas of functional space in need of attention owing to negative effects on ecosystem function. For example, the loss of functionally unique large frugivores from the San Antonio assemblage likely increased the potential for collapse of mutualistic networks of bird and plant species in which previous work revealed they play a critical role (Palacio et al. 2016). Loss of large frugivores can further have negative cascading effects on ecosystem health by reducing seed dispersal, affecting the survival of native trees, influencing the potential for forest regeneration, and even constraining the vegetation’s ability to respond to climate change (Cordeiro & Howe 2003; Moran et al. 2009; McConkey et al. 2012; Mokany et al. 2014; Ribeiro da Silva et al. 2015; Bovo et al. 2018).

Although the San Antonio bird assemblage has changed in functional volume, the core of its functional fingerprint (concentrating a majority of species which are arguably functionally redundant) has retained its position. High functional redundancy may be just as important as richness extending to the periphery of functional space for ecosystem health, being one of the mechanisms helping to maintain high diversity and ecosystem function (Wohl et al. 2004), reducing negative effects of natural enemies of species (Philpot et al. 2012), and facilitating niche packing within assemblages (Ricklefs 2012; Pigot et al. 2016; Cooke et al. 2019a). Our finding that the core of the San Antonio bird assemblage has seemingly not lost functions nor shifted in position suggests that there are still important attributes of the system’s functionality that mirror the pre-disturbance assemblage of 100+ years ago. Hopefully, given the right conditions (i.e. time, continued forest recovery and increased connectivity between remaining forest fragments), San Antonio may still recover more of its lost functionality provided by extirpated species, some of which are are still found in the wider region (Palacio et al. 2016, 2019).

Our study provides the first example of an assessment of temporal changes in funcional fingerprints from a highly diverse tropical ecosystem, and both our findings and the methods we employed have wide applications particularly for tropical ecosystem study and conservation (Stroud & Thompson 2019). For instance, restoration objectives aimed at replicating previous states of a system, could use measures of functional fingerprints to quantify how much change is needed and to direct efforts to areas of functional space in particular need of recovery (Meli et al. 2014). Similarly, conservation actions aimed at maintaining or recovering functionality can use functional fingerprints to assess routes to recovery and to prioritize species with a particular combination of traits which should be preserved or potentially re-introduced (Chazdon 2008). Future studies could explore whether the San Antonio functional fingerprints can be used as a basis for comparison to other tropical localities lacking long-term data (Stroud & Thompson 2019), and other historical suveys (e.g. Miller et al. 1957) can be used to carry out comparisons over larger scales.

Certain species traits, such as being large, specialized, social, having low dispersal ability and being at the top end of food webs, make animal species more vulnerable to local extirpation (Kattan et al. 1994; Davies et al. 2000; Pearson et al. 2009; Habel et al. 2019). Our results highlight that some of these traits, namely body size, dispersal ability and habitat breadth, are responsible for ~78% of the variation in functional diversity within a tropical bird assemblage. This means that species with trait values at the extremes of these axes of variation (size, dispersal and habitat specialization) likely also account for a large proportion of the functional richness and dispersion in ecological assemblages. Thus, when a system loses species at the extremes of these axes, it likely also loses unique functionality. In San Antonio, the extirpation of species combining some of these traits associated with vulnerability resulted in significant decreases in functional richness and dispersion, hence arguably making the assemblage less healthy and potentially more susceptible to further changes (Mouillot et al. 2013). In agreement with our results, species vulnerability to extinction is positively correlated to functional uniqueness and specialization in a wide range of organisms (Pimiento et al. 2020), and the functions provided by these unique species may be particularly prone to disappear (Mouillot et al. 2013; Teichert et al. 2017). Therefore, conservation efforts aimed at maintaining ecosystem health must move beyond just maintaining species numbers to designing strategies for the maintenance of ecological function by identifying and conserving species with traits conferring high vulnerability (Cadotte et al. 2011; McConkey et al. 2012; Pimiento et al. 2020).

## Acknowledgements

We dedicate this paper to the memory of Gustavo H. Kattan, who spearheaded studies of changes in bird assemblages in San Antonio and contributed enormously to Neotropical ecology and conservation through his career. Because of Gustavo’s untimely passing we were unable to share this work with him and to invite him to contribute as a coauthor.

J. Von Rothkirch, M. L. Mahecha and Y. Caicedo contributed with specimen measurements; A. M. Cuervo, Glenn Zeeholzer, Amanda Rodewald’s Lab group and Daniel Cadena’s Lab group provided excellent feedback; The following scientific collections allowed access to bird specimens: IAvH, ICN, Museo de La Salle, Museo Univalle, AMNH.

## Supporting information

Appendix S1 contains the complete trait dataset used in this tudy plus the PC scores and the values of functional Uniqueness and Distinctiveness estimated for each species. Appendix S2 contains the R code used for all analyses, and Appendix S3 contains supplementary methods, tables and figures.

## Appendix S3. Supplementary methods, figures and tables

### Functional trait dataset

Our trait dataset contains a combination of measurements taken from almost 1900 bird specimens housed at seven ornithological collections in Colombia, compiled mainly by Montoya et al. (2018) and complemented with secondary information from other published trait datasets (Claramunt et al. 2012; Cooke et al. 2019; Pigot et al. 2020). For 93 species which did not have complete information, we measured additional specimens (see acknowledgements). Where possible, at least 2 males and 2 females of each species were measured (mean = 8, range = 1 - 124 individuals). Habitat breadth (the number of habitats listed in IUCN accounts) and generation time (years), were extracted from the IUCN List of Threatened Species accounts (BirdLife International 2018). Mean trait values for each species were estimated and used in all subsequent analyses. The impact of measurement error due to multiple observers on our analysis is negligable due to the large disparity in body sizes in our dataset (eg. *Coragyps atratus* ‘~1,600 g vs. *Ocreatus underwoodii* ~2,5 g).

### Historical changes in the San Antonio Avifauna

The original avifauna, which contained 201 species in the 1910’s, suffered the extirpation of 47 species between the 1930’s and 1990’s due to extensive forest fragmentation and hunting (Kattan et al. 1994). However, 17 species were reestablished in the 2000’s after forest regeneration (Palacio et al. 2019). Additionally, since the 1990’s the site has been colonized by 32 species not recorded in the 1910’s (Kattan et al. 1994; Palacio et al. 2019). Large canopy frugivores, understory insectivores, habitat specialists and species at their elevational limits were more prone to extirpation in San Antonio (Kattan et al. 1994), in agreement with work in other tropical areas experiencing forest fragmentation (Renjifo 1999; Sodhi et al. 2006; Ferraz et al. 2007; Pearson et al. 2009; Stouffer 2020).

**Supplementary Table S3.1.** Summary of PCA showing standard deviation, variance explained and loadings for 9 PC axes. 79% of the variation in functional traits was explained by the first 3 PC. PC1 describes body size, PC2 dispersal ability and PC3 habitat breadth.

|  | **PC1** | **PC2** | **PC3** | **PC4** | **PC5** | **PC6** | **PC7** | **PC8** | **PC9** |
| --- | --- | --- | --- | --- | --- | --- | --- | --- | --- |
| Standard deviation | 2.23 | 1.09 | 0.98 | 0.90 | 0.64 | 0.54 | 0.45 | 0.35 | 0.28 |
| Proportion of variance | 0.55 | 0.13 | 0.11 | 0.09 | 0.05 | 0.03 | 0.02 | 0.01 | 0.01 |
| Cumulative proportion | 0.55 | 0.68 | 0.79 | 0.88 | 0.92 | 0.96 | 0.98 | 0.99 | 1.00 |
| **Loadings** | **PC1** | **PC2** | **PC3** | **PC4** | **PC5** | **PC6** | **PC7** | **PC8** | **PC9** |
| Weight | 0.41 | 0.12 |  | 0.22 |  | 0.11 | 0.58 | 0.20 | 0.62 |
| Bill length | 0.33 |  |  | -0.66 |  | 0.44 | -0.32 | 0.37 | 0.12 |
| Bill width | 0.36 | 0.17 | -0.19 | -0.48 |  | -0.25 | 0.41 | -0.53 | -0.24 |
| Wing length | 0.41 | -0.14 | 0.12 | 0.22 |  |  | 0.22 | 0.47 | -0.69 |
| Tail length | 0.37 |  |  | 0.17 | -0.71 | -0.41 | -0.37 |  | 0.16 |
| Tarsus | 0.37 | 0.18 |  | 0.39 | 0.14 | 0.56 | -0.28 | -0.49 | -0.11 |
| HWI |  | **-0.85** | 0.31 | -0.15 | -0.19 | 0.20 | 0.13 | -0.26 |  |
| Habitat breadth |  | -0.37 | **-0.89** | 0.20 |  |  |  |  |  |
| Generation time | 0.36 | -0.22 | 0.17 |  | 0.65 | -0.46 | -0.35 |  | 0.16 |

**Figure S3.1.**
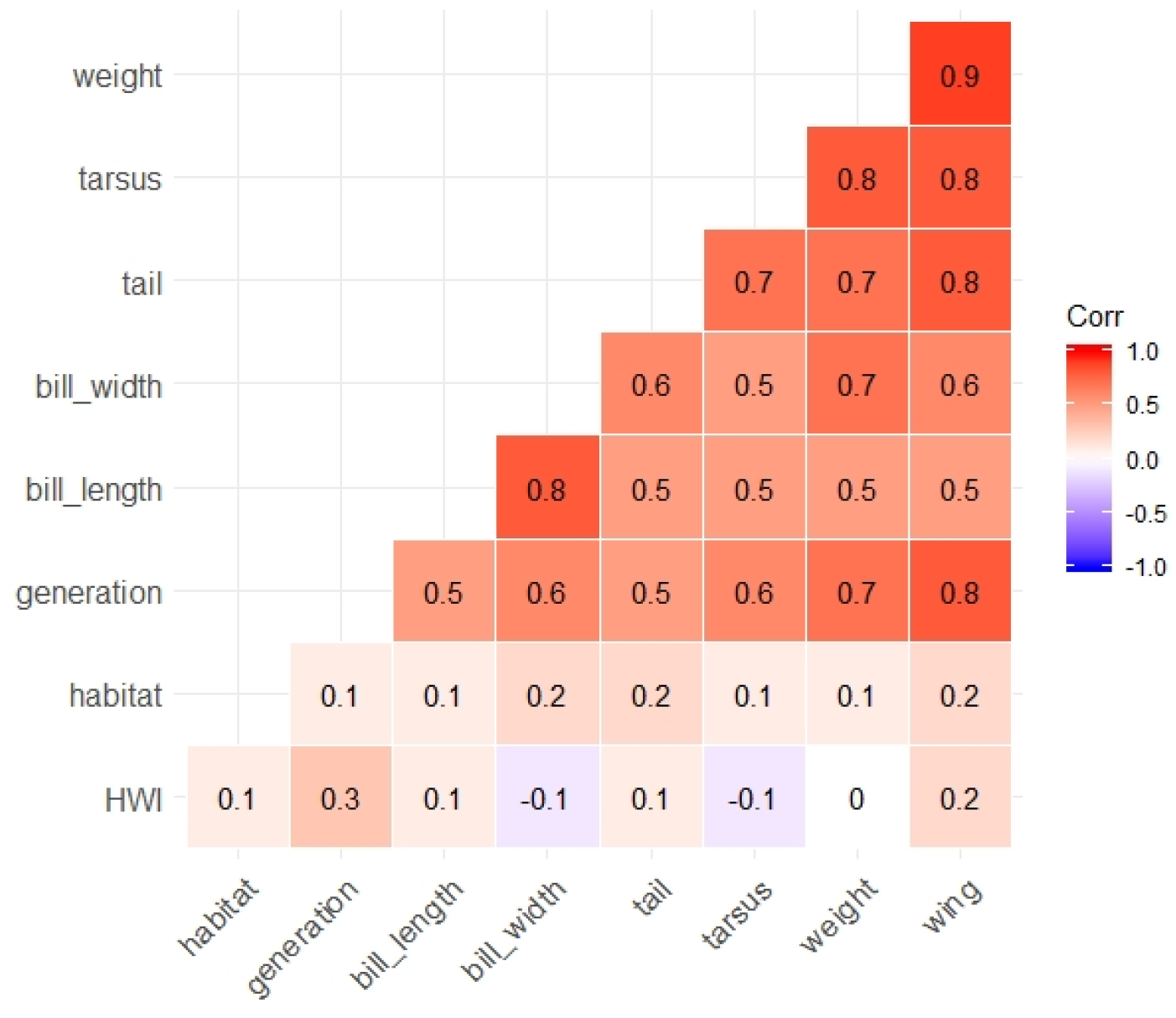
Correlation matrix of the 9 functional traits used in this study. 72% of our traits have correlation values equal to or below ±0.6.

**Figure S3.2.**
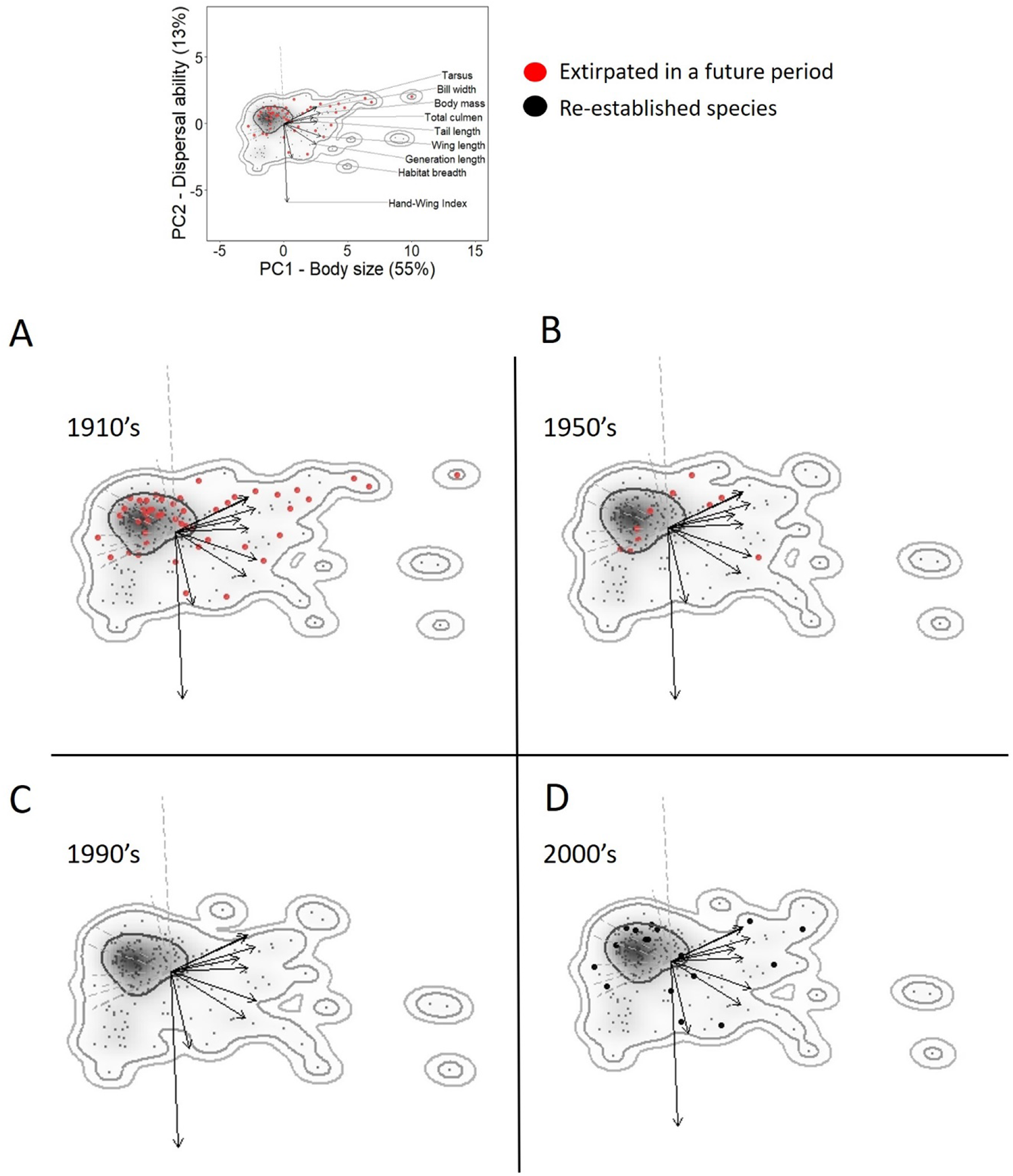
Species extirpations and re-establishments (Scenario 2) have caused changes in the shape and extent of the functional space of San Antonio in 100 years, but the centroid of trait space (gray shading) has not shifted. Two-dimensional functional space is represented by PC scores of functional traits during 4 time periods (**A-D**). PC1 reflects largely variation in body size while PC2 correlates with dispersal ability of birds and habitat breadth. Small dots are species present in each time period, with larger red dots representing species which became extirpated in a future period and larger black dots are species which re-established populations after being extirpated. Arrows are scaled to represent the loadings and direction of each trait in functional space and the insert above shows the scale and the traits represented by each arrow. Gray shading represents the kernel density estimates for each time period and curved lines show the 50, 95 and 99% probability contours.

**Figure S3.3.**
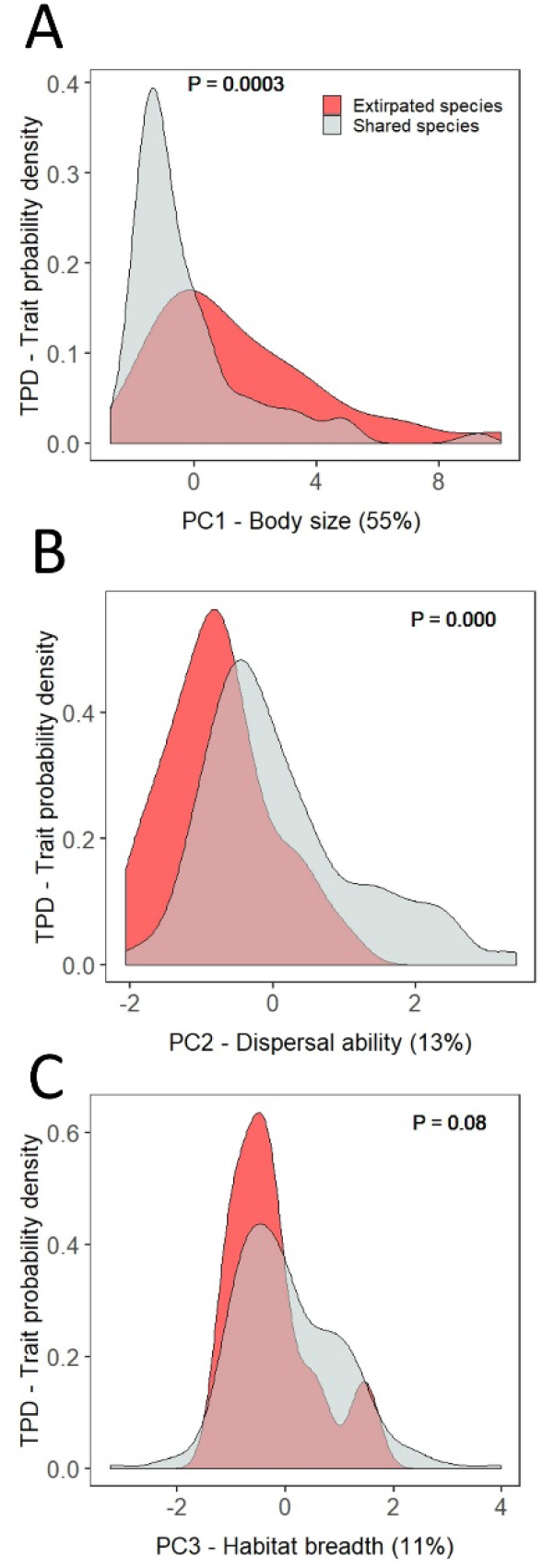
Under Scenario 2, assuming there were no novel species joining the San Antonio bird assemblage, the probability density curves of extirpated species between 1910’s and 2000’s, show that the functional fingerprint has shifted towards **A.** Smaller birds, and **B.** Species with higher dispersal ability. **C.** Contrasting with scenario 1, in scenario 2 species habitat breadths did not change significantly as a consequence of species extirpations. *P* values were obtained through Kolmogorov-Smirnov tests comparing the distributions.

